## Supplementary material for "Replay as wavefronts and theta sequences as bump oscillations in a grid cell attractor network": Figure supplements

### Figure supplements for "Replay as wavefronts and theta sequences as bump oscillations in a grid cell attractor network"

Louis Kang[1] and Michael R. DeWeese[1]

[1] *Redwood Center for Theoretical Neuroscience, Helen Wills Neuroscience Institute, and Department of Physics, University of California, Berkeley, Berkeley, California, USA*

(Dated: October 25, 2019)

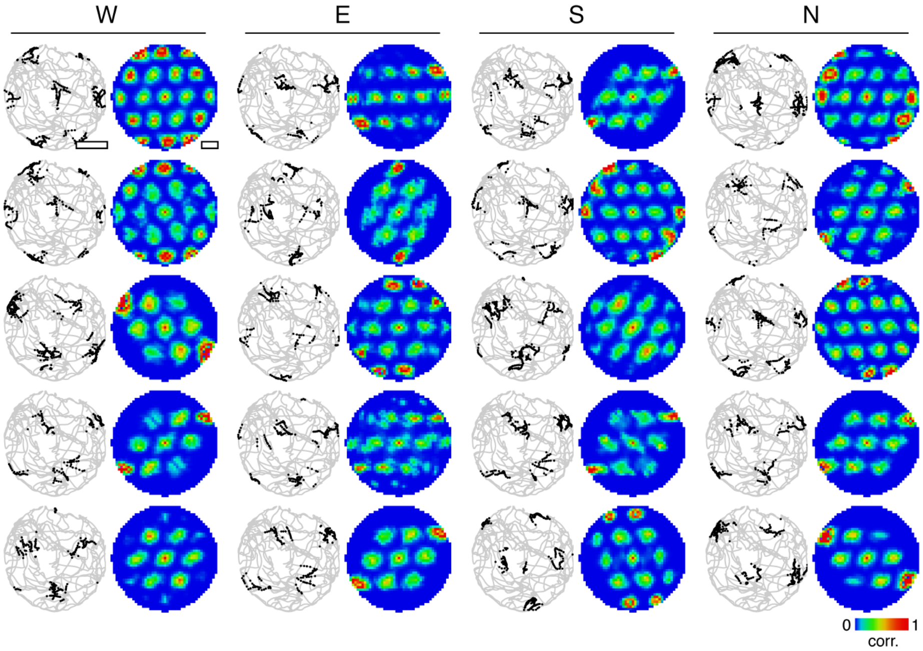


**Figure 1–figure supplement 1**. Spatial firing maps for a 2D open field trajectory. Twenty grid cells are selected from the simulation depicted in **Figure 1**D,E across all four excitatory populations. For each grid cell, spikes superimposed on the animal's trajectory (left) and autocorrelation of rate maps calculated from spikes (right).

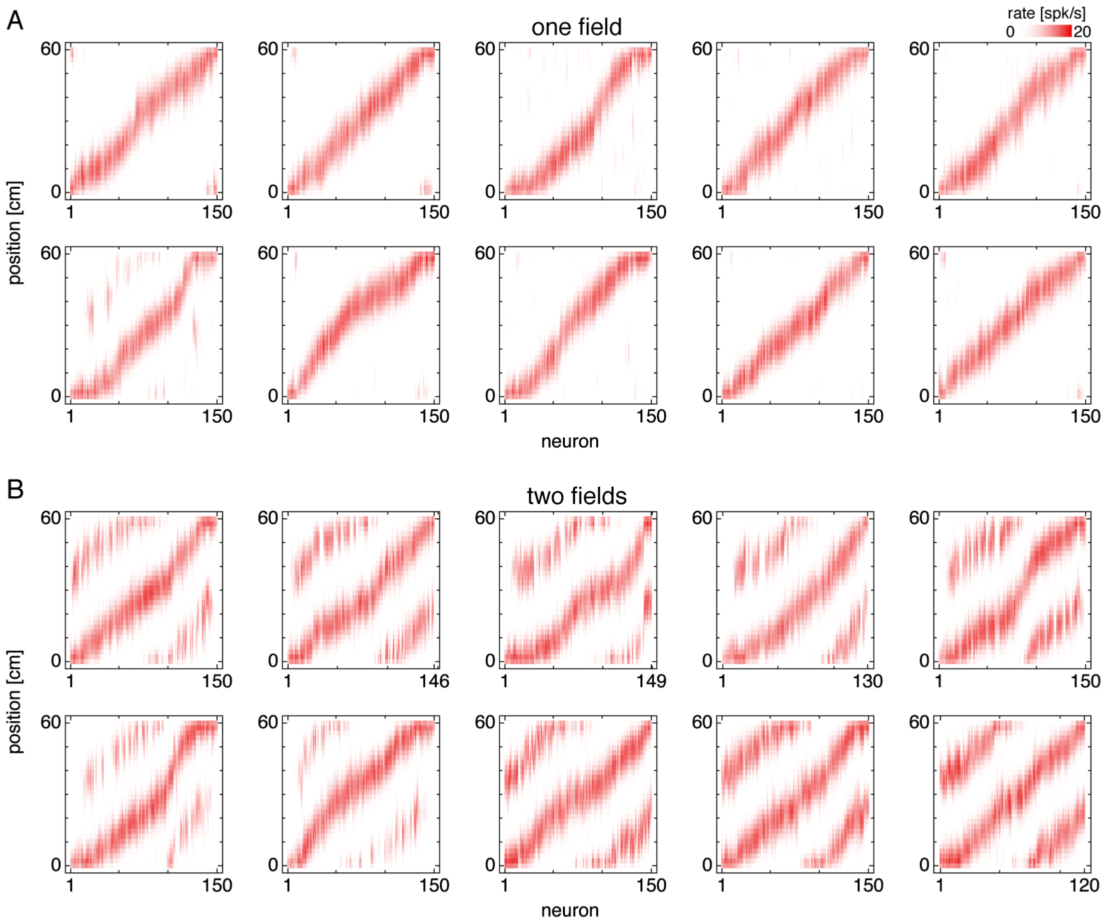


**Figure 2–figure supplement 1**. Firing fields of recorded grid cells along a 1D track. Fields are sorted by position of maximum rate. (**A**) Representative simulations exhibiting one field. (**B**) Representative simulations exhibiting two fields.

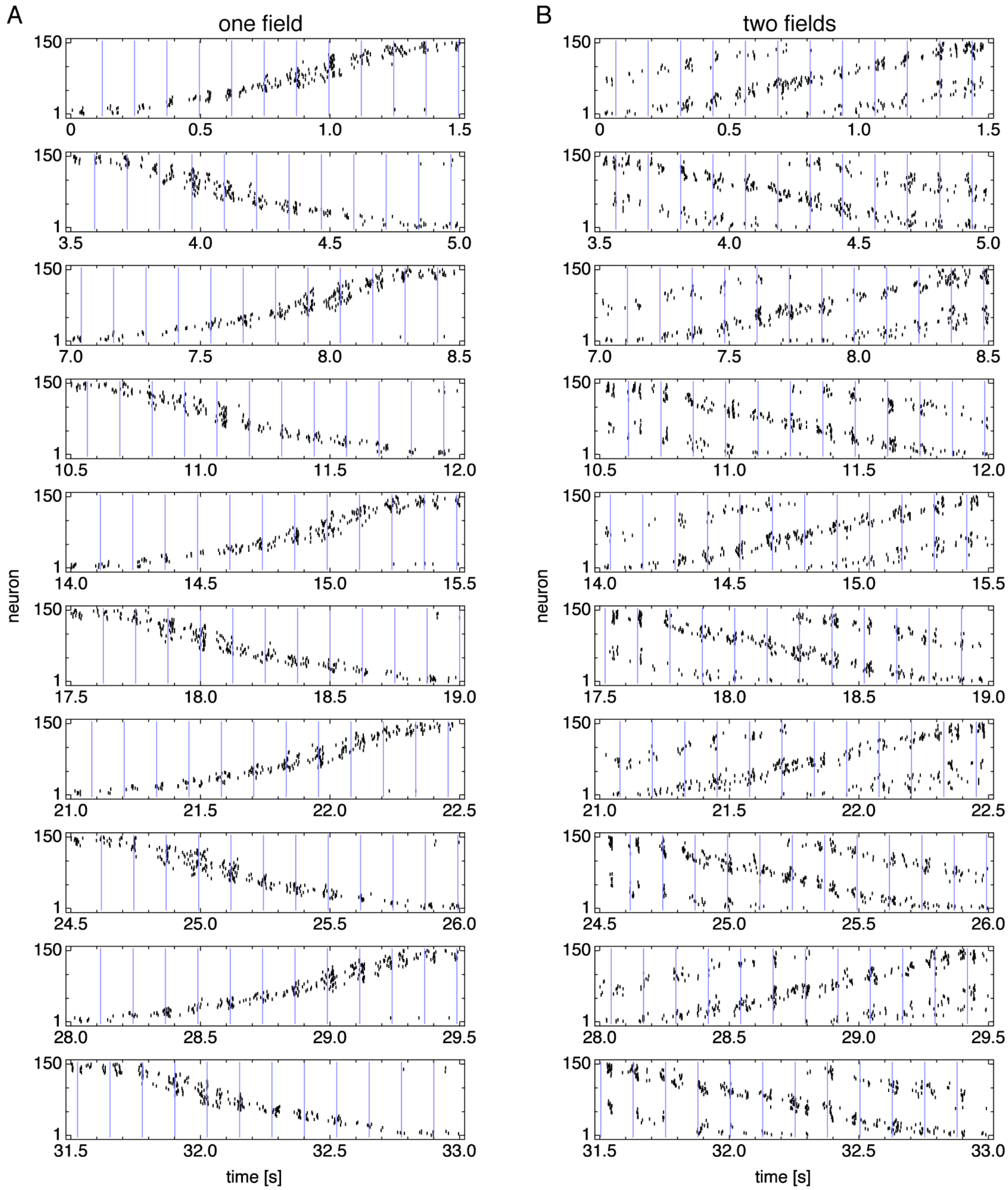


**Figure 3–figure supplement 1**. Spike rasters over multiple runs. Vertical blue lines indicate theta cycle boundaries. (**A**) Representative simulation exhibiting one field. (**B**) Representative simulation exhibiting two fields.

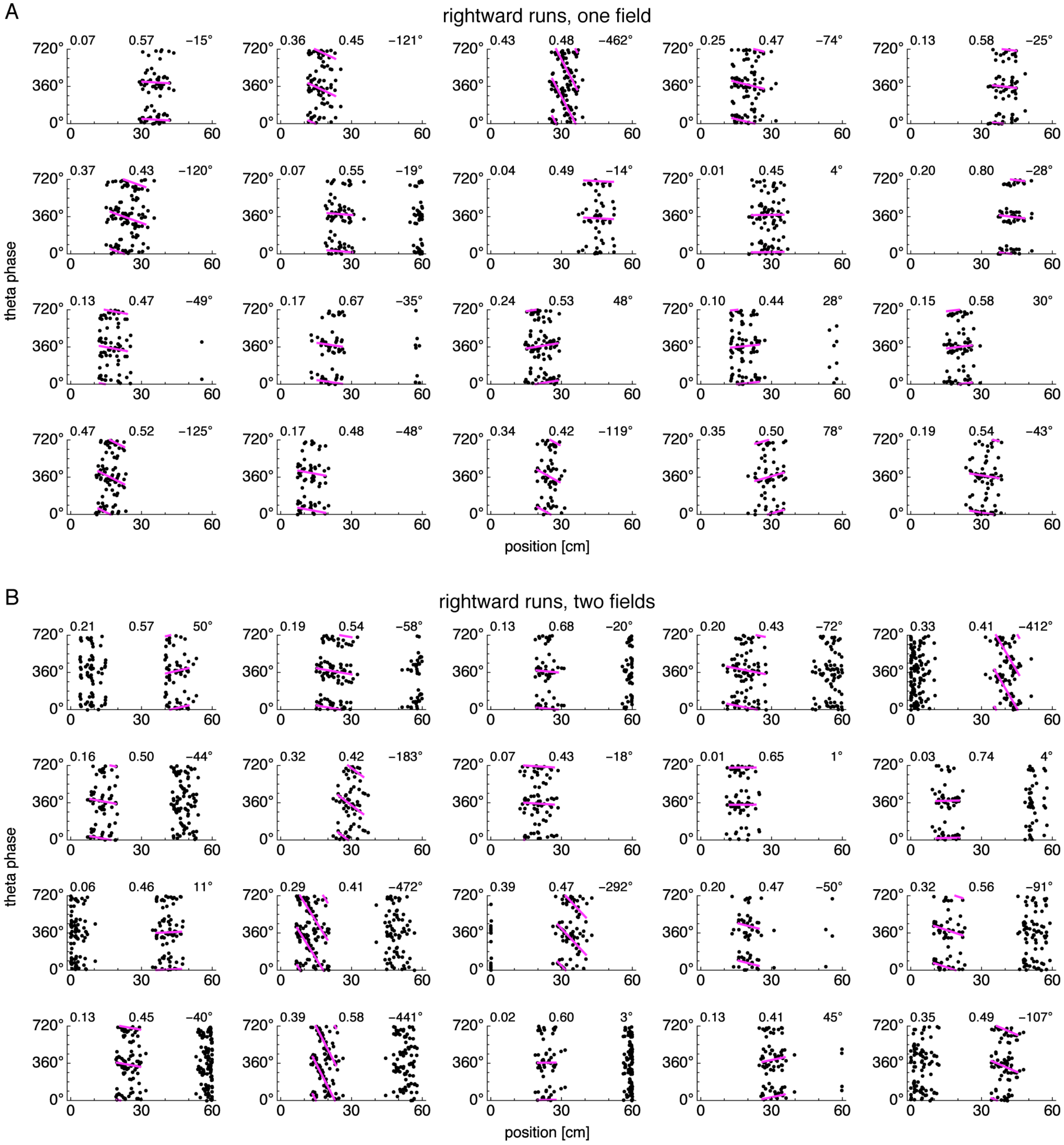


**Figure 3–figure supplement 2**. Relationship between animal position along the track and theta phase during rightward runs. Dots represent spikes during runs in the directions indicated by arrows, with each spike repeated at two equivalent phases for clarity. Lines indicate fit by circular-linear regression. Numbers in each panel from top left to top right indicate magnitudes of correlation coefficients, regression fit scores, and precession ranges. (**A**) Representative neurons from simulations exhibiting one field. (**B**) Representative neurons from simulations exhibiting two fields.

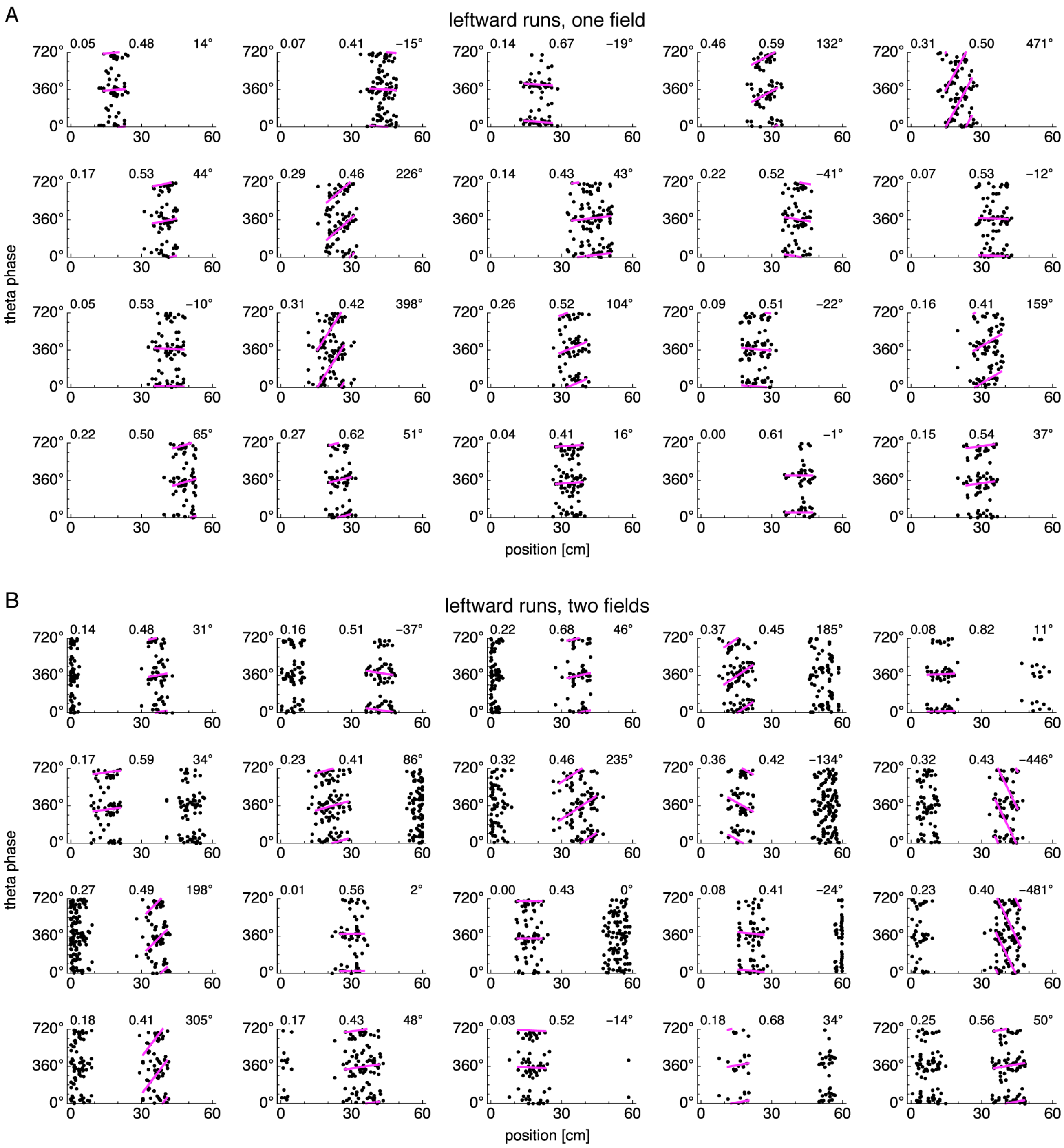


**Figure 3–figure supplement 3**. Relationship between animal position along the track and theta phase during leftward runs. Panels same as in Figure 3–figure supplement 2.

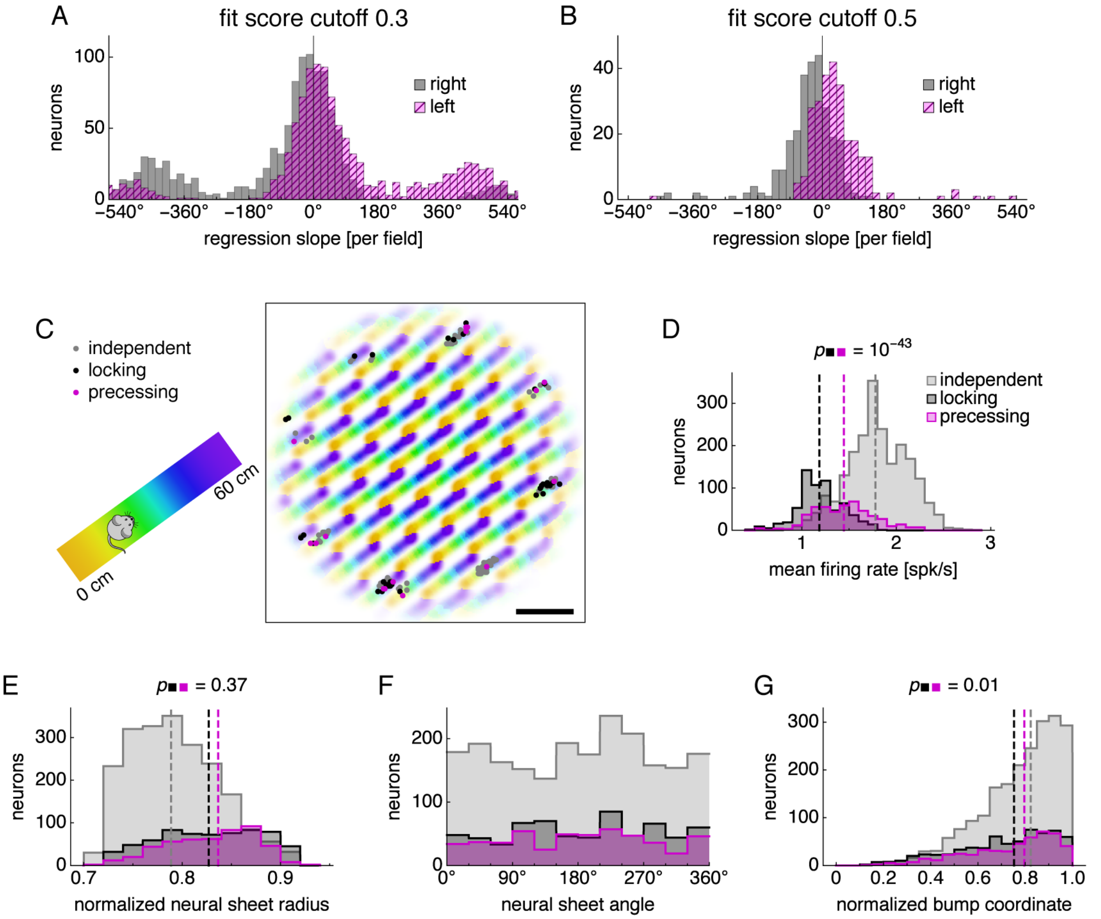


**Figure 3–figure supplement 4**. Further analysis of subgroups of phase-independent, phase-locking, and phase-precessing neurons. (**A,B**) Distributions of regression slopes using a fit score cutoff different from 0.4, which was used in **Figure 3**. (**C**) Neural sheet locations of neurons belonging to each subgroup for the simulation described in **Figure 3**A,B. Data encompassing recording areas shown in **Figure 2**C and recording areas offset by 45° on the neural sheet. Left, track diagram. Right, neural activity over runs with each neuron colored according to the track position at which it attained maximum firing rate. Scale bar, 50 neurons. (**D–G**) Distribution of neurons in each subgroup with respect to various properties. Medians for each subgroup indicated by dashed lines. Medians for phase-locking and phase-precessing subgroups compared by the Mann-Whitney $U$ test with reported $p$-value. (**D**) Mean firing rate. (**E**) Distance from the center of the neural sheet divided by the half the neural sheet size $n$. (**F**) Angle on the neural sheet relative to the right direction, or $+\hat{\mathbf{x}}$. (**G**) Location relative to attractor bumps in units of relative firing rate. A value of 1 means that the center of a bump passes over the neuron, and a value of 0 means that no part of a bump passes over the neuron (**Appendix A**).

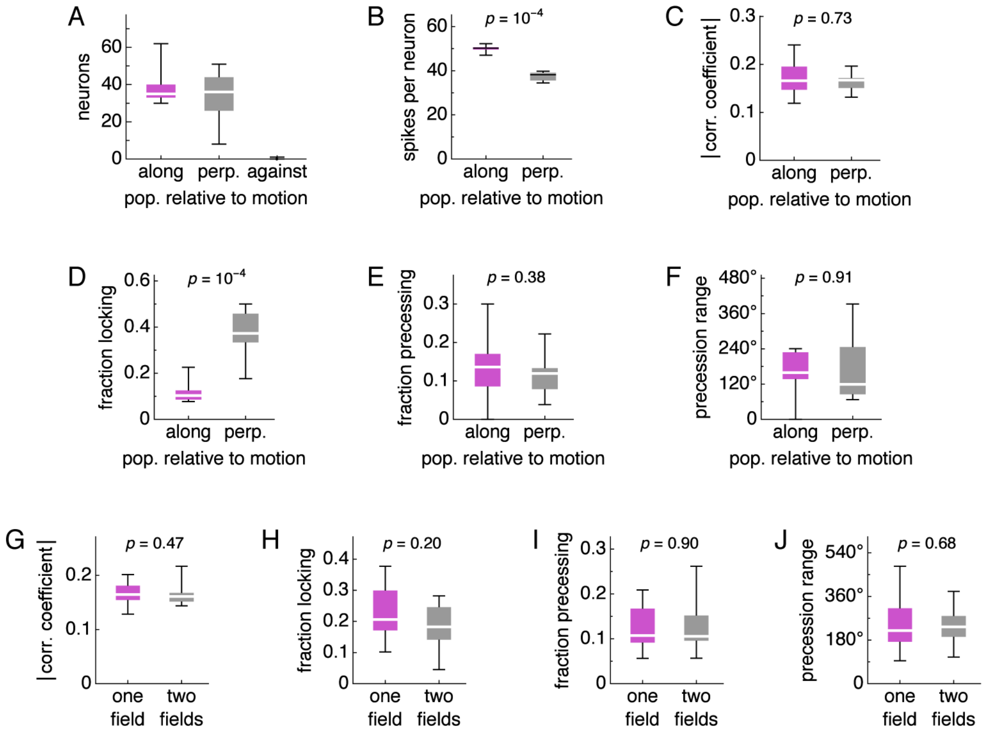


**Figure 3–figure supplement 5**. Box-whisker plots with lines at medians, boxes between first to third quartiles, and whiskers between the entire data range excluding outliers lying more than 1.5 times the interquartile distance beyond the first or third quartile. Medians compared by the Mann-Whitney $U$ test with reported $p$-values. (**A**–**F**) Analysis of phase relationships with respect to excitatory population. For these simulations, the direction of animal velocity is 0°, which means that runs are aligned to the E and W directions. The "along" category corresponds to both the E population when the animal is running towards E and the W population when the animal is running towards W. The "against" category corresponds to the E and W populations during the opposite running directions. The "perp." category corresponds to the N and S directions. (**A**) Number of valid recorded neurons. Note that there are very few "against" neurons due to low firing rate, and this category is excluded from further analysis. (**B**) Number of spikes per neuron. (**C**) Magnitude of correlation coefficient. (**D**) Fraction of phase locking neurons. (**E**) Fraction of phase precessing neurons. (**F**) Phase precession range. (**G**–**J**) Analysis of phase relationships with respect to number of fields exhibited by a simulation. (**G**) Magnitude of correlation coefficient. (**H**) Fraction of phase locking neurons. (**I**) Fraction of phase precessing neurons. (**J**) Phase precession range.

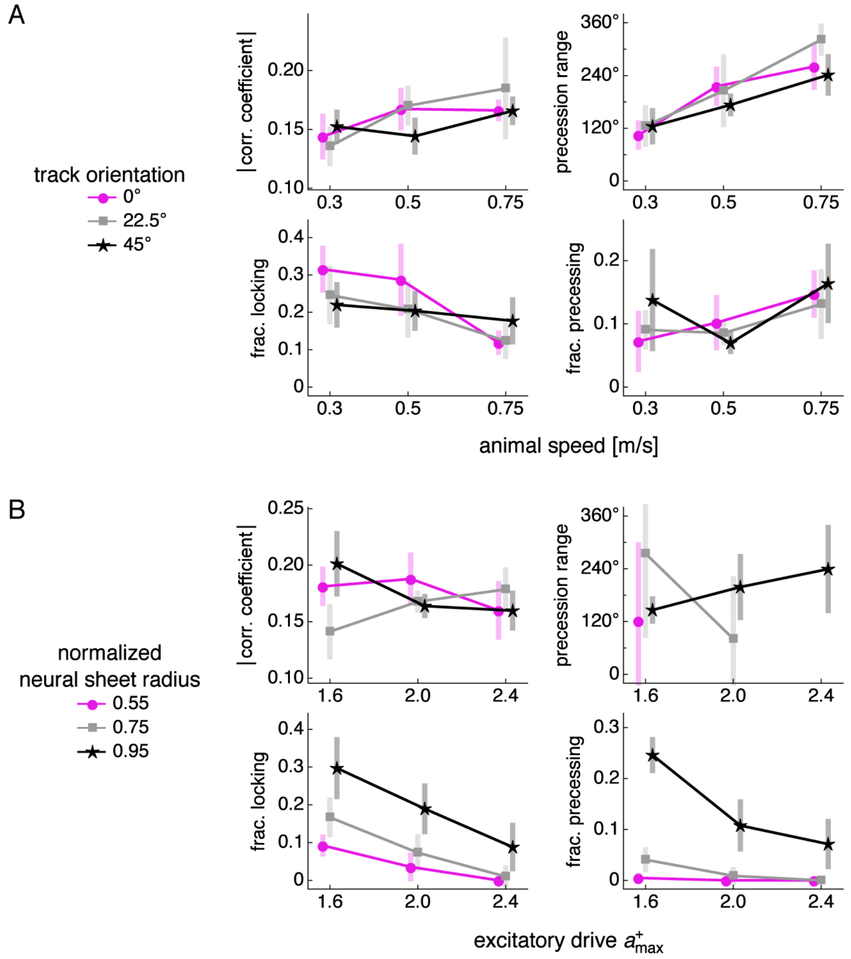


**Figure 3–figure supplement 6**. Correlation coefficient magnitude, precession range, fraction of phase-locking neurons, and fraction of precessing neurons as functions of various simulation parameters. Points at mean values with bars indicating s.d. over replicate simulations. Small horizontal offsets added for clarity. (**A**) Animal speed and track orientation relative to the E direction, or $+\hat{\mathbf{X}}$. (**B**) Excitatory drive and recording distance from the center of the neural sheet divided by the half the neural sheet size.

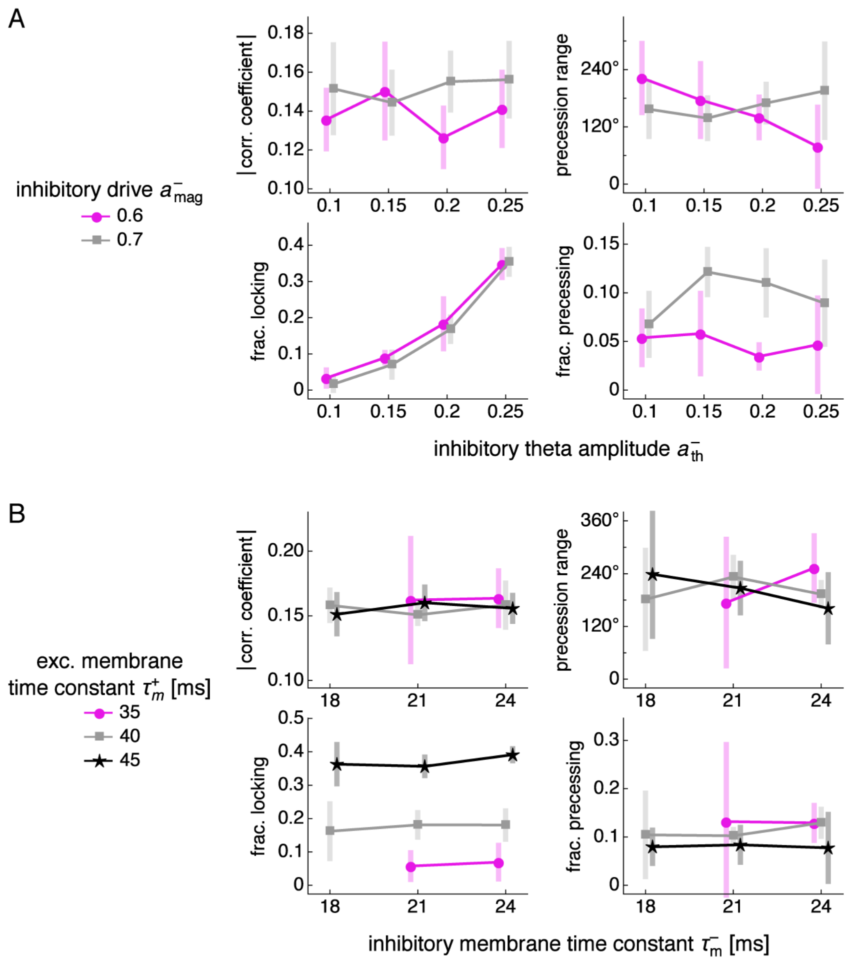


**Figure 3–figure supplement 7**. Same as Figure 3–figure supplement 6, except as functions of different parameters. (**A**) Inhibitory theta amplitude and inhibitory drive. (**B**) Inhibitory and excitatory membrane time constants.

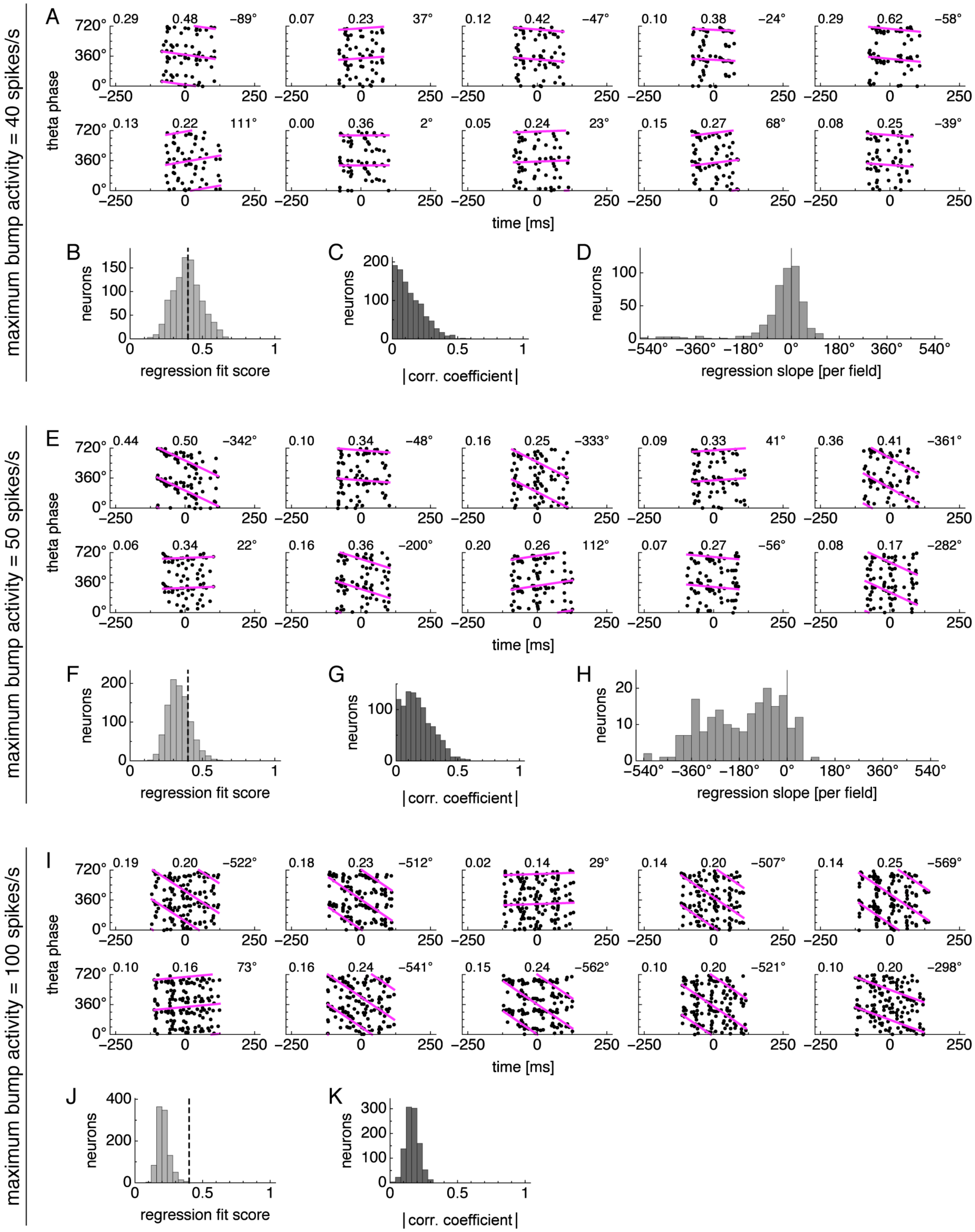


**Figure 4–figure supplement 1**. Simplified model of phase precession using average bump dynamics. Bump activity is rescaled to different maximum values. (**A**–**D**) Neurons exhibit phase locking under low bump activity with maximum value 40 spikes/s. Statistics from 1000 neurons simulated according to **Figure 4**E. (**A**) Relationship between animal position along the track and theta phase for representative neurons. Dots represent spikes. Lines indicate fit by circular-linear regression. Numbers in each panel from top left to top right indicate magnitude of correlation coefficient, regression fit score, and regression slope. (**B**) Magnitudes of circular-linear correlation coefficients. (**C**) Fit scores for circular-linear regression. (**D**) Regression slopes for neurons with fit score >0.4. (**E**–**H**) Neurons exhibit phase precession under medium bump activity with maximum value 50 spikes/s. Panels same as **A**–**D**. (**I**–**K**) Neurons exhibit phase independence under high bump activity with maximum value 100 spikes/s. Panels same as **A**–**C**.

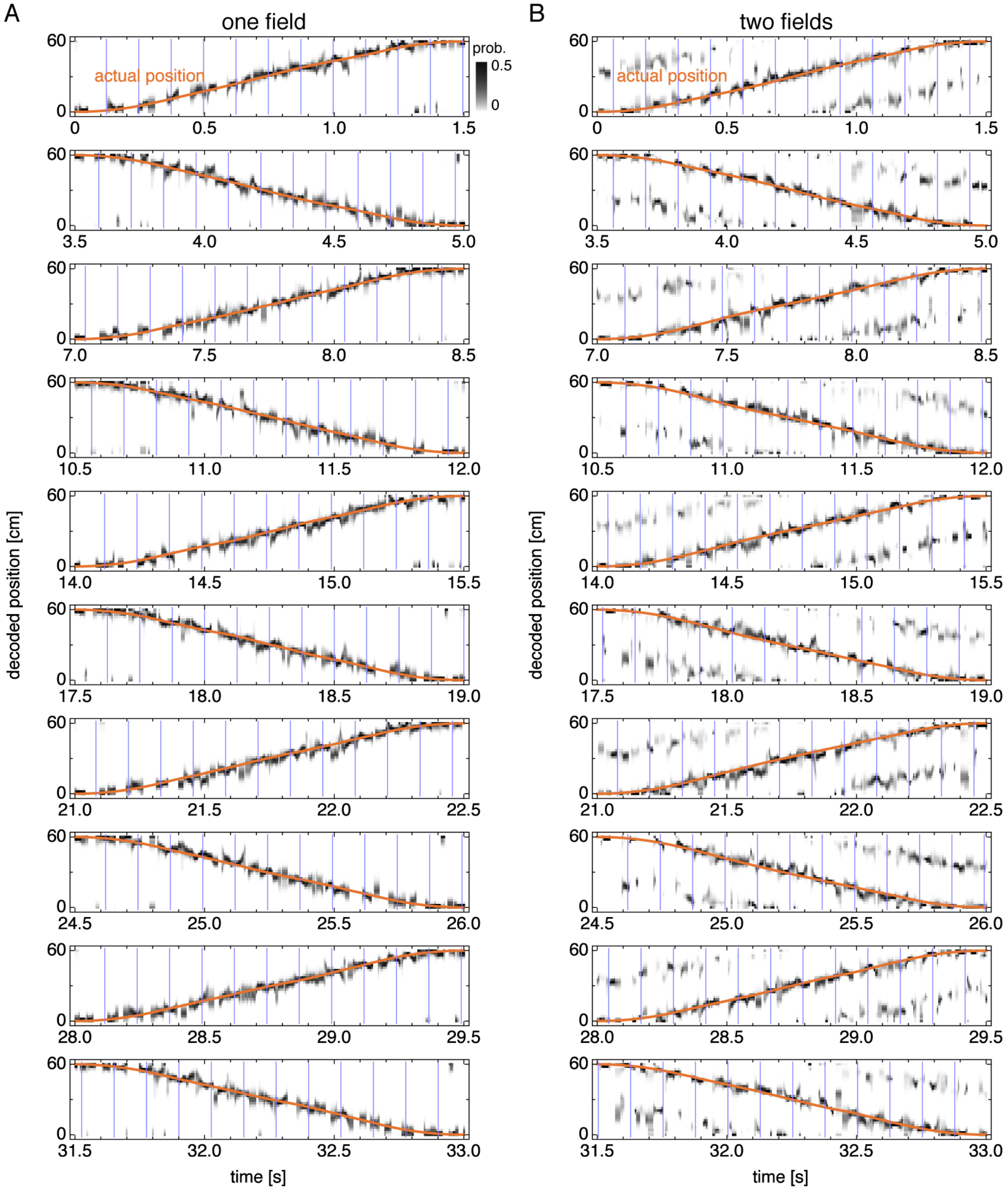


**Figure 5–figure supplement 1**. Decoded position over multiple runs corresponding to Figure 3–figure supplement 1. Vertical blue lines indicate theta cycle boundaries. Orange line indicates the actual animal trajectory. (**A**) Representative simulation exhibiting one field. (**B**) Representative simulation exhibiting two fields.

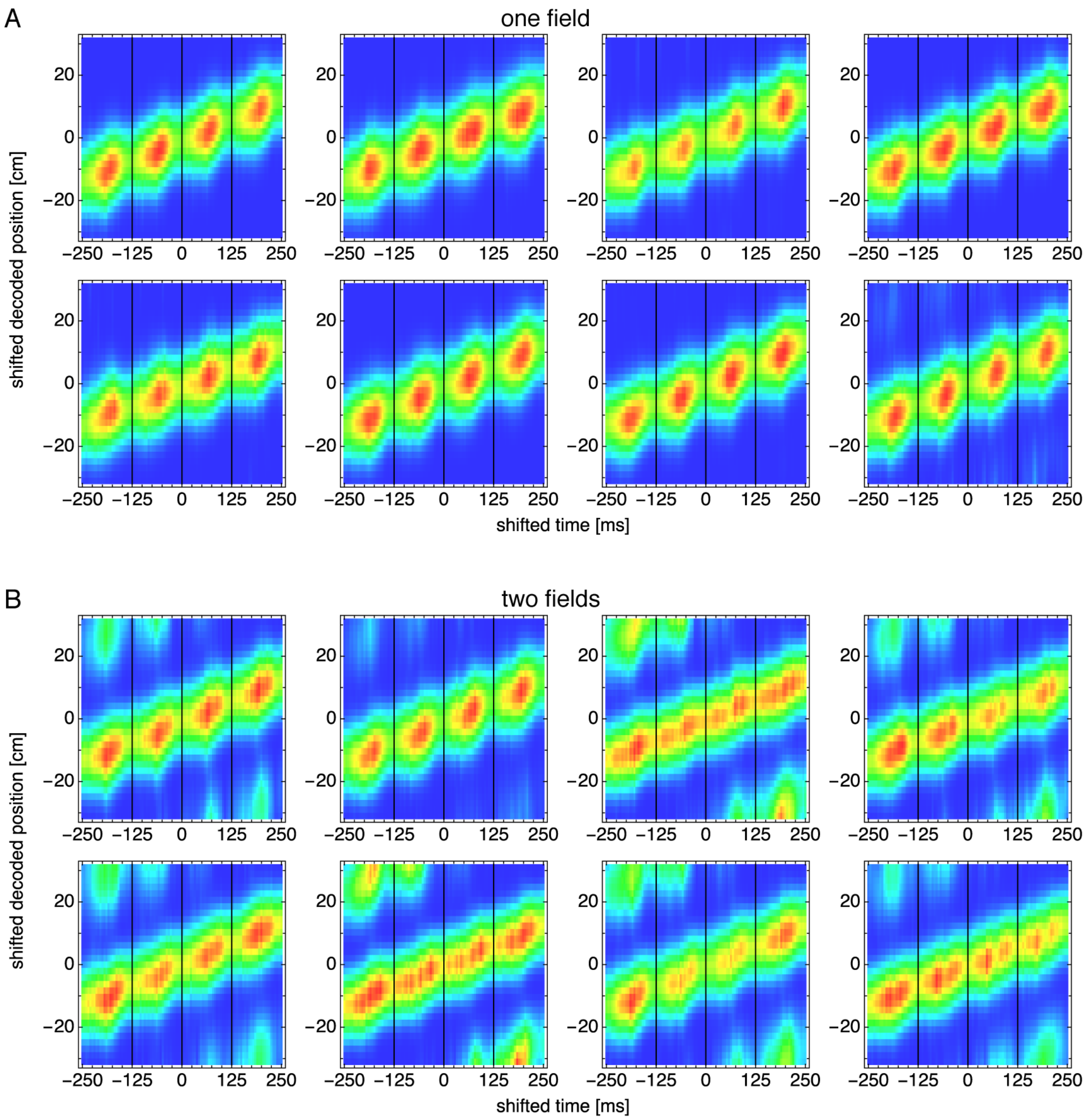


**Figure 5–figure supplement 2**. Decoded position shifted by actual position and averaged over theta cycles for many simulations. Vertical black lines indicate theta cycle boundaries. (**A**) Representative simulations exhibiting one field. (**B**) Representative simulations exhibiting two fields.

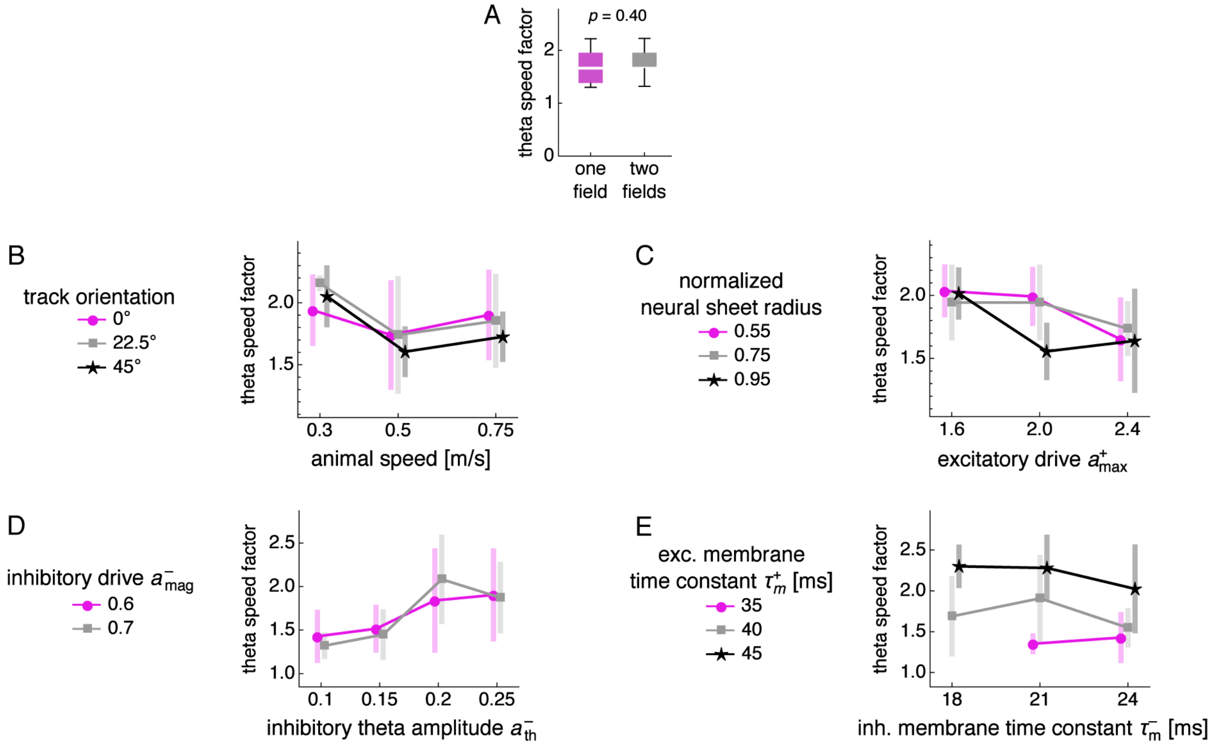


**Figure 5–figure supplement 3**. Theta sequence speed divided by mean actual speed for different numbers of fields and parameter values. (**A**) Dependence of theta speed on number of fields. Box-whisker plot with line at medians, box between first to third quartiles, and whiskers between the entire data range excluding outliers lying more than 1.5 times the interquartile distance beyond the first or third quartile. $p$-value calculated by the Mann-Whitney $U$ test. (**B**–**E**) Theta sequence speed divided by mean actual speed as a function of various simulation parameters. Points at mean values with bars indicating s.d. over replicate simulations. (**B**) Animal speed and track orientation relative to the E direction, or $+\hat{\mathbf{X}}$. (**C**) Excitatory drive and recording distance from the center of the neural sheet divided by the half the neural sheet size. (**D**) Inhibitory theta amplitude and inhibitory drive. (**E**) Inhibitory and excitatory membrane time constants.

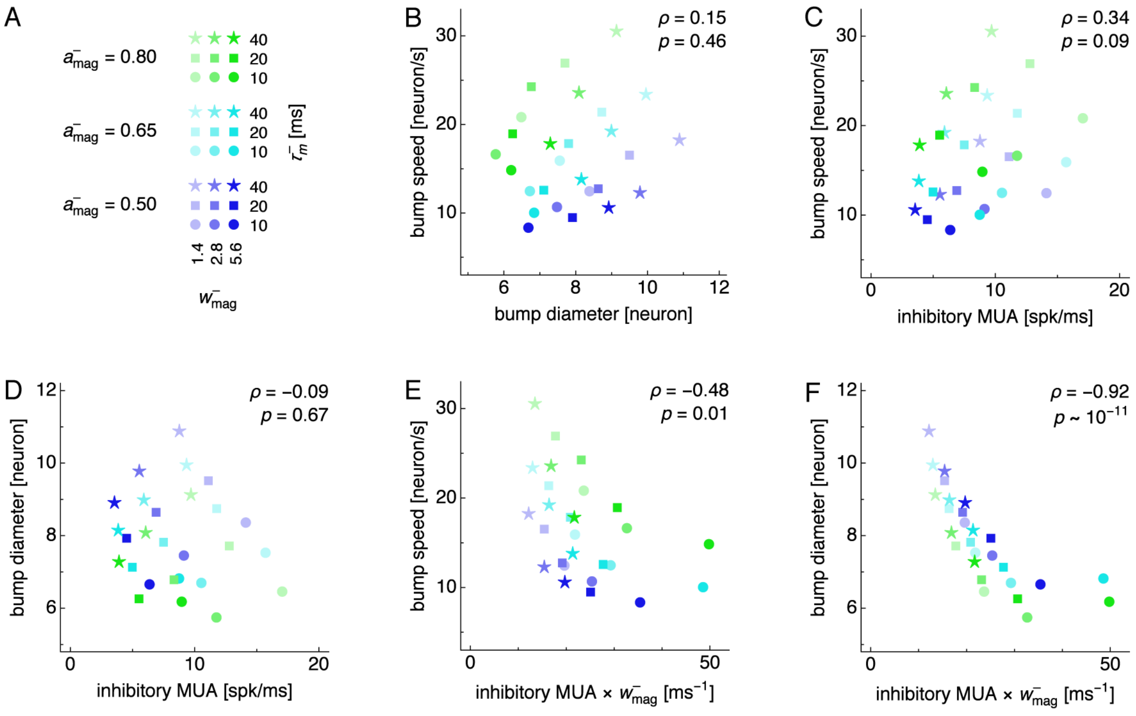


**Figure 6–figure supplement 1**. Attractor bump size and speed are not generally correlated over changes in multiple simulation parameters, but both are negatively correlated with the amount of inhibition produced by inhibitory neurons. (**A**) Legend for simulations with fixed inhibitory drive. Each simulation is performed at different values of inhibitory drive $a^-_{\mathrm{mag}}$, inhibitory synaptic strength $w^-_{\mathrm{mag}}$, and inhibitory membrane time constant $\tau^-_{\mathrm{m}}$. Other parameters are set at the values used for main simulation runs (**Video 1**). (**B–F**) Relationships between pairs of attractor bump properties across parameters displayed in **A**. Rank correlation computed by Spearman's $\rho$ with $p$-values indicated. Attractor bump properties are diameter, speed, the multiunit activity of inhibitory neurons (inhibitory MUA), and a metric for the amount of inhibition produced by inhibitory neurons (inhibitory MUA $\times\, w^-_{\mathrm{mag}}$). Note that **B** indicates no overall correlation between bump size and speed, but increasing $a^-_{\mathrm{mag}}$ (following blue–cyan–green) increases bump speed and decreases bump diameter, as seen in **Figures 4**B and **6**B.

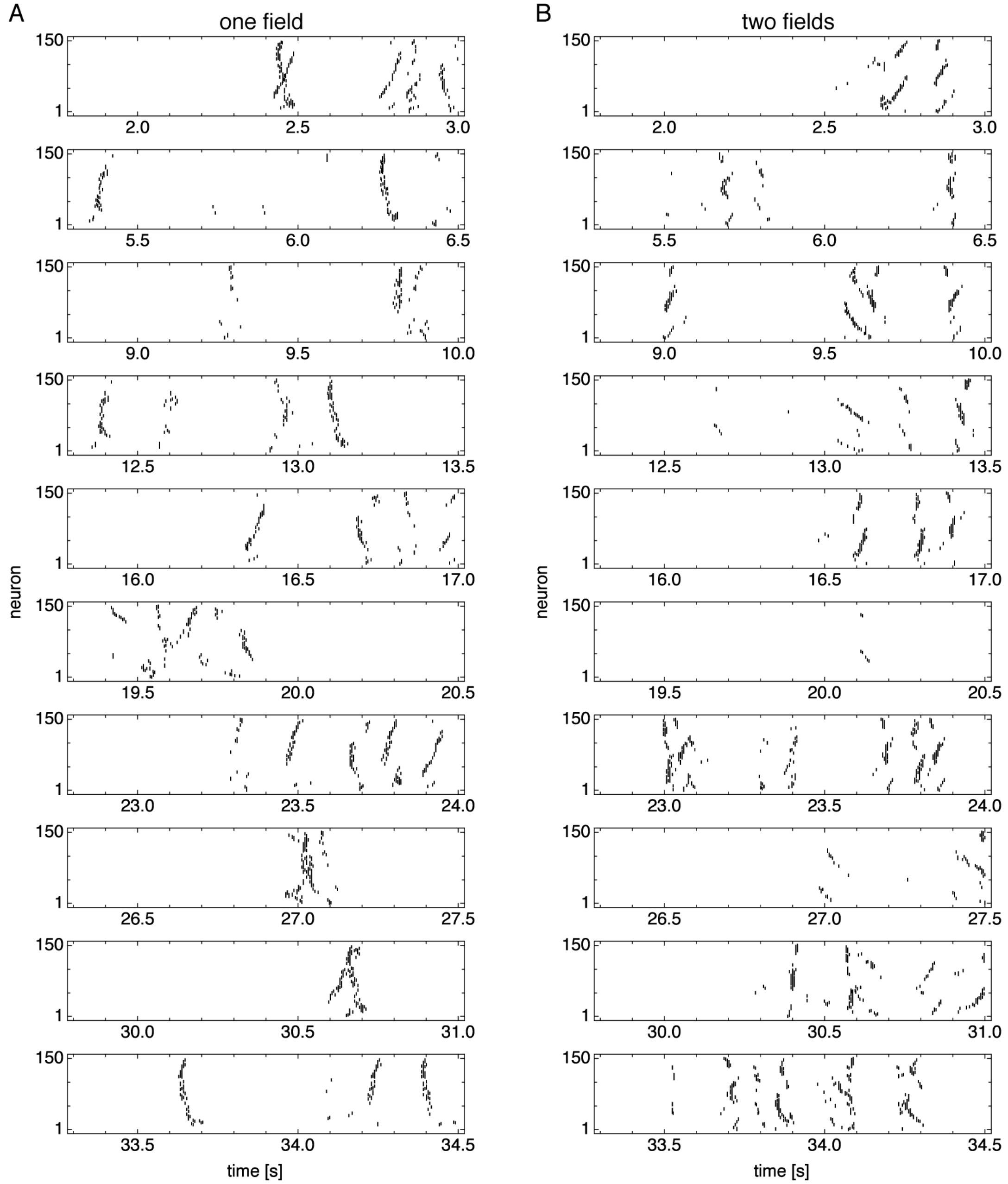


**Figure 7–figure supplement 1**. Spike rasters over multiple idle periods. (**A**) Representative simulation exhibiting one field. (**B**) Representative simulation exhibiting two fields.

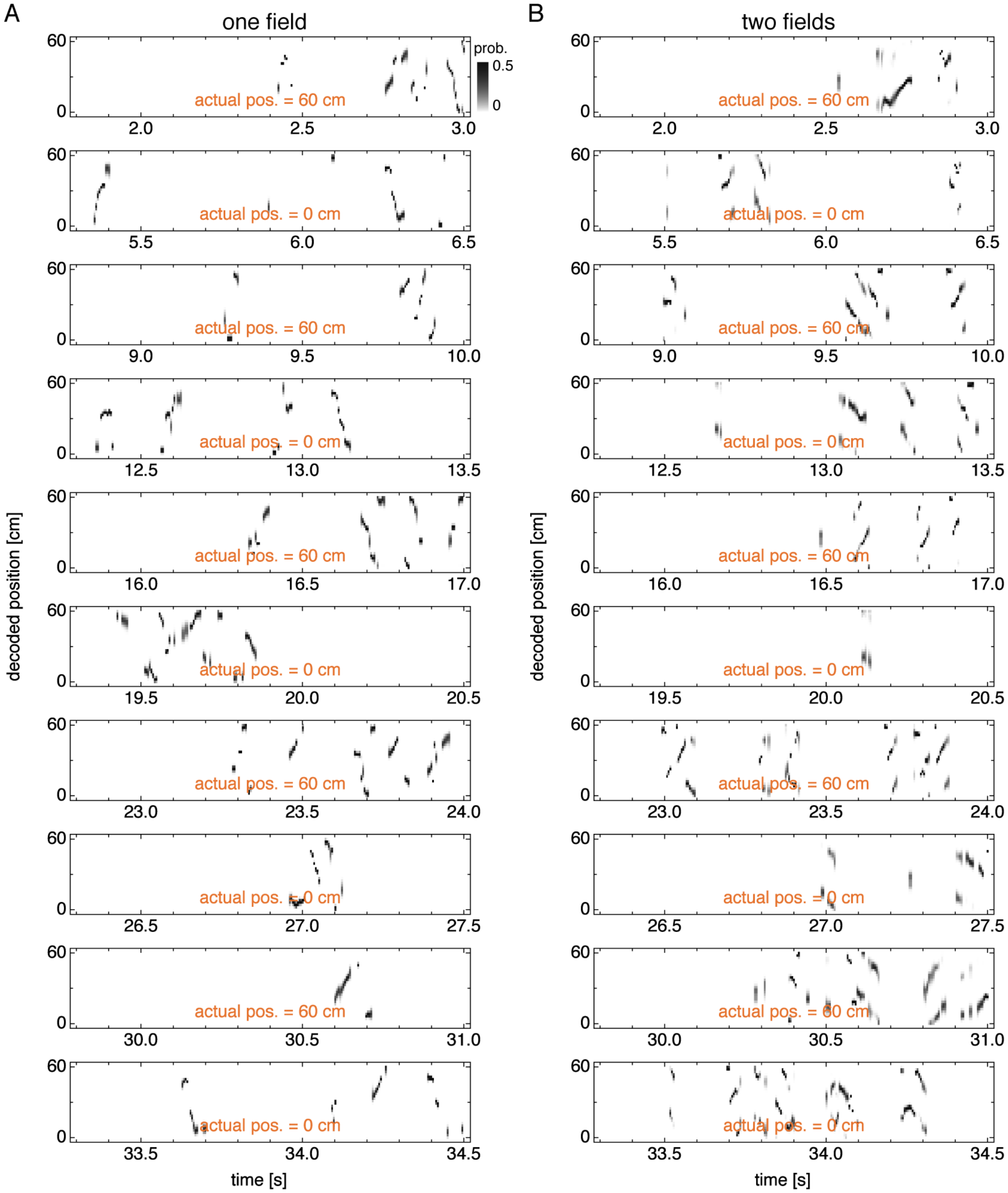


**Figure 7–figure supplement 2**. Decoded position over multiple idle periods corresponding to Figure 7–figure supplement 1. (**A**) Representative simulation exhibiting one field. (**B**) Representative simulation exhibiting two fields.

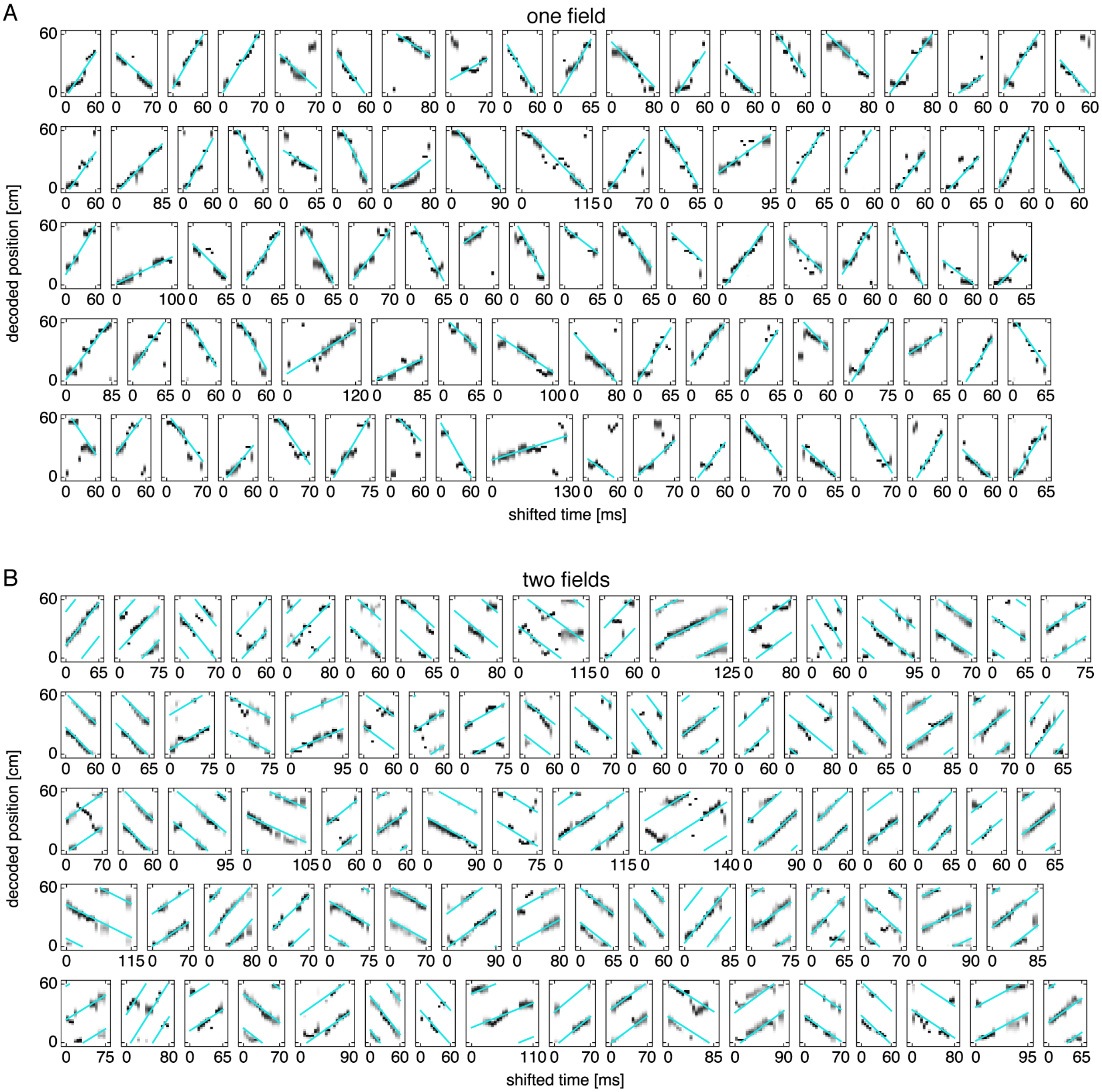


**Figure 7–figure supplement 3**. Replays with cyan fit lines. (**A**) Representative replays from simulations exhibiting one field. (**B**) Representative replays from simulations exhibiting two fields. Note that two or three parallel lines participate in the fit (**Appendix A**).

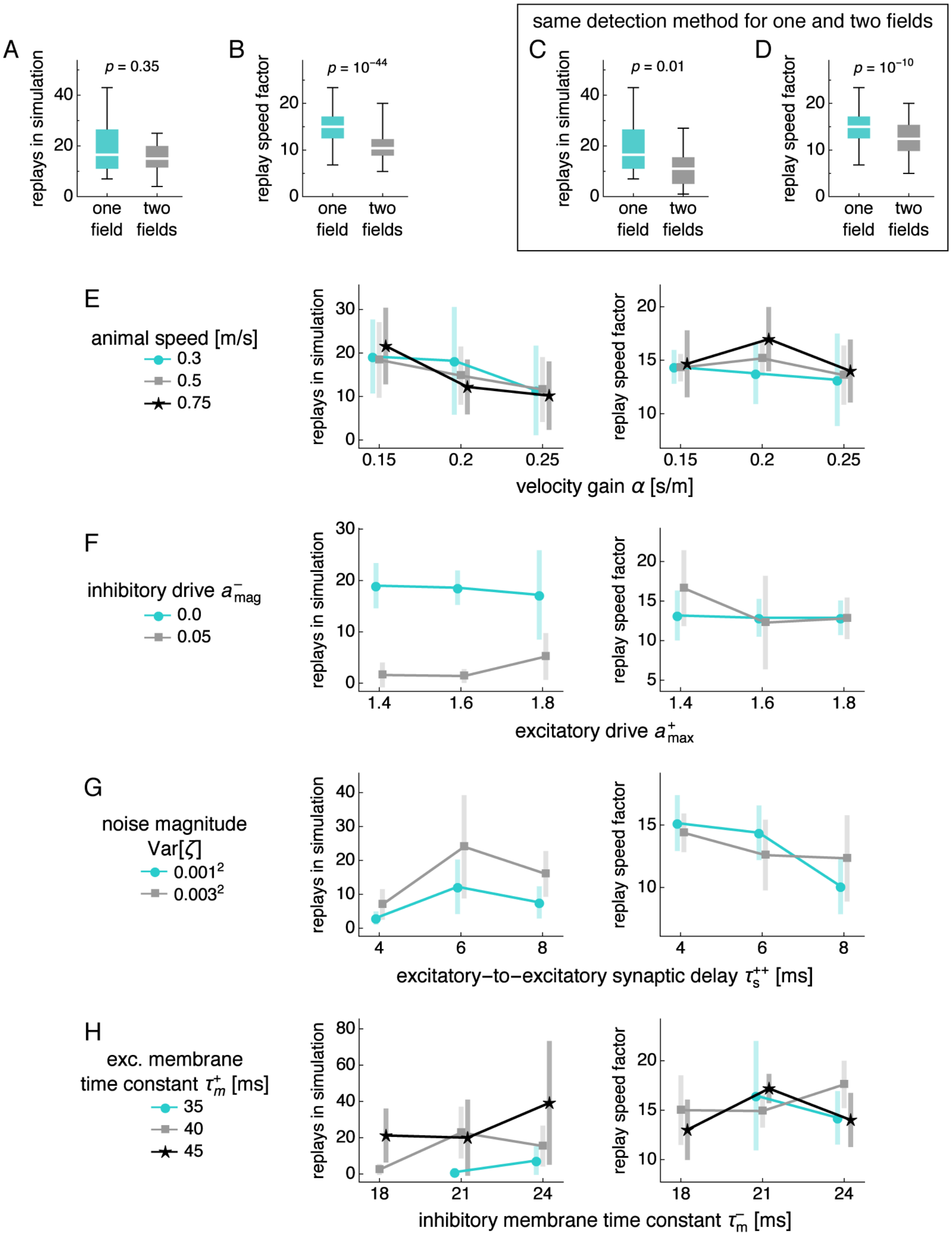


**Figure 7–figure supplement 4.** (**A–D**) Dependence of replay count and speed on number of fields. Box-whisker plot with line at medians, box between first to third quartiles, and whiskers between the entire data range excluding outliers lying more than 1.5 times the interquartile distance beyond the first or third quartile. $p$-value calculated by the Mann-Whitney $U$ test. (**A**) Replay count. (**B**) Replay speed divided by mean actual speed. (**C,D**) Same as **A,B**, but we detect replays in simulations exhibiting two fields with the same method used for single fields. (**E–H**) Replay count and replay speed divided by mean actual speed as functions of various simulation parameters. Points at mean values with bars indicating s.d. over replicate simulations. (**E**) Velocity gain and animal speed. (**F**) Excitatory and inhibitory drive. (**F**) Excitatory-to-excitatory synaptic delay and noise magnitude for both excitatory and inhibitory neurons. (**H**) Inhibitory and excitatory membrane time constants.

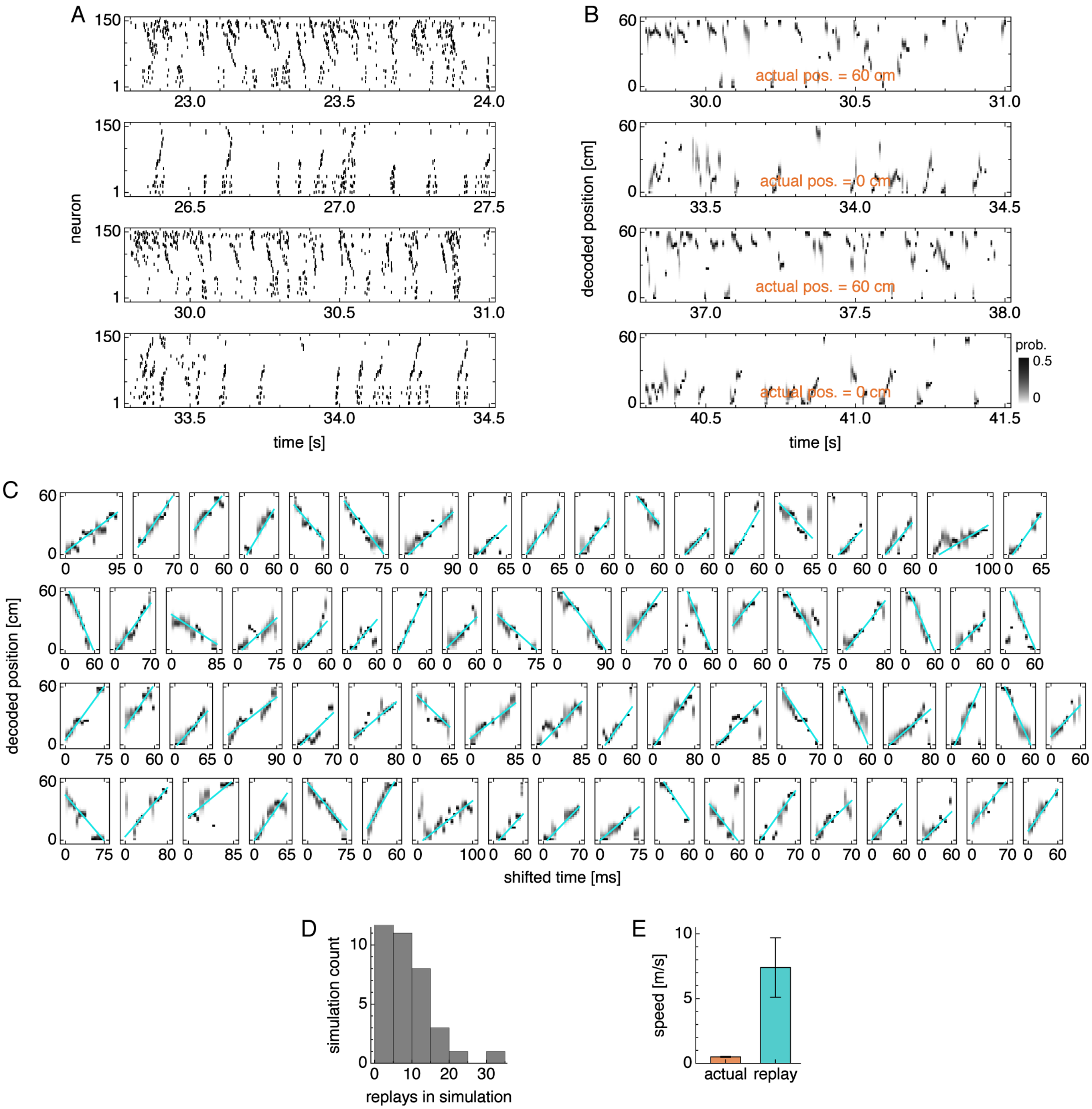


**Figure 8–figure supplement 1**. Replay properties in simulations with lower velocity gain. (**A**) Grid cell spike rasters for four idle periods. (**B**) Decoded position corresponding to **A**. (**C**) Replays across multiple simulations with cyan fit lines. (**D**) Number of replays across simulations. (**E**) Actual run speed $0.50 \pm 0.05\,\mathrm{m/s}$ (mean $\pm$ s.d. over time) and replay speed $7.4 \pm 2.3\,\mathrm{m/s}$ (mean $\pm$ s.d. over replays).

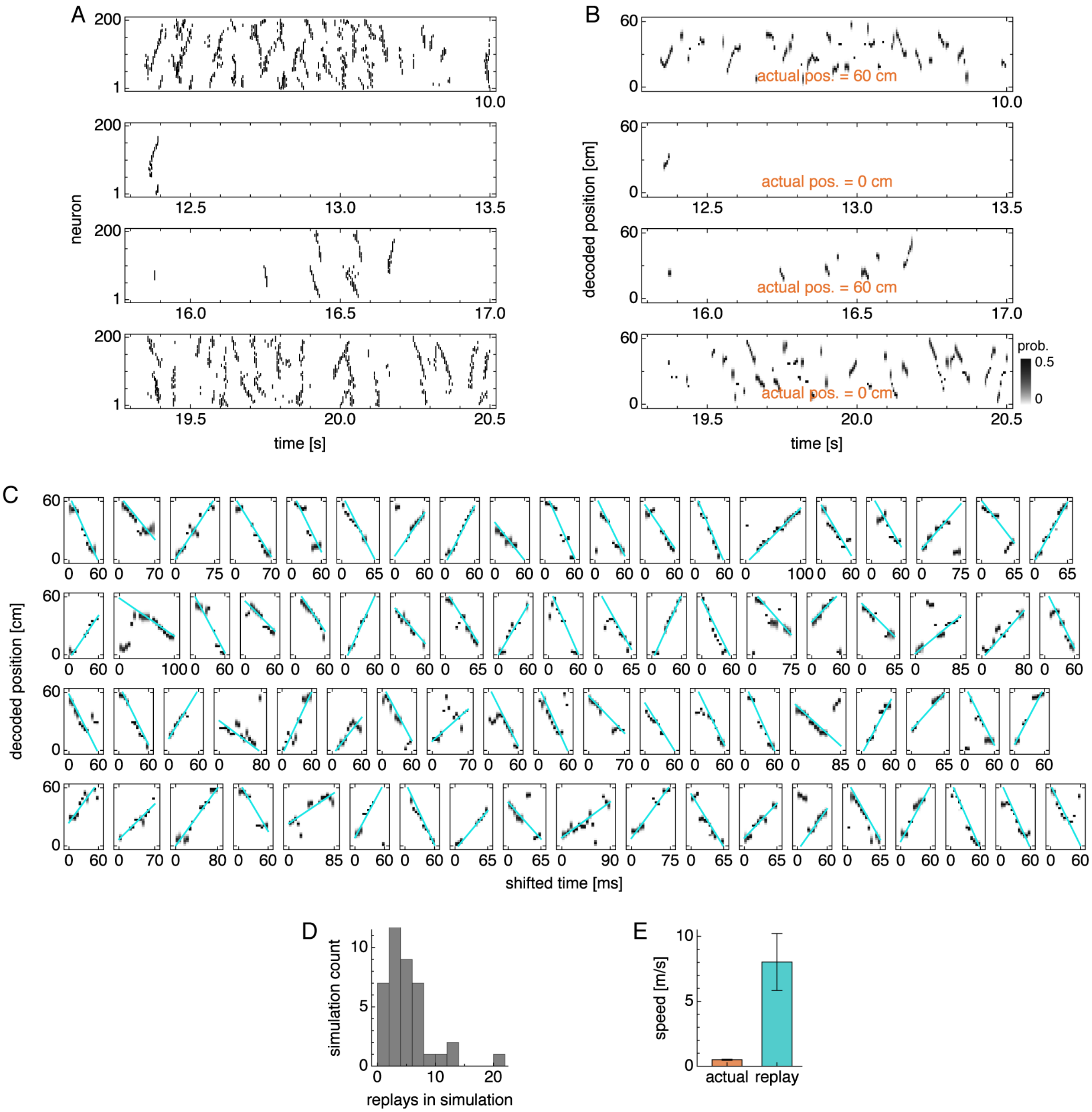


**Figure 8–figure supplement 2**. Same as Figure 8–figure supplement 1, but for simulations with periodic boundary conditions and uniform excitatory drive. Replay speed is $8.0 \pm 2.2\,\mathrm{m/s}$ (mean $\pm$ s.d. over replays).

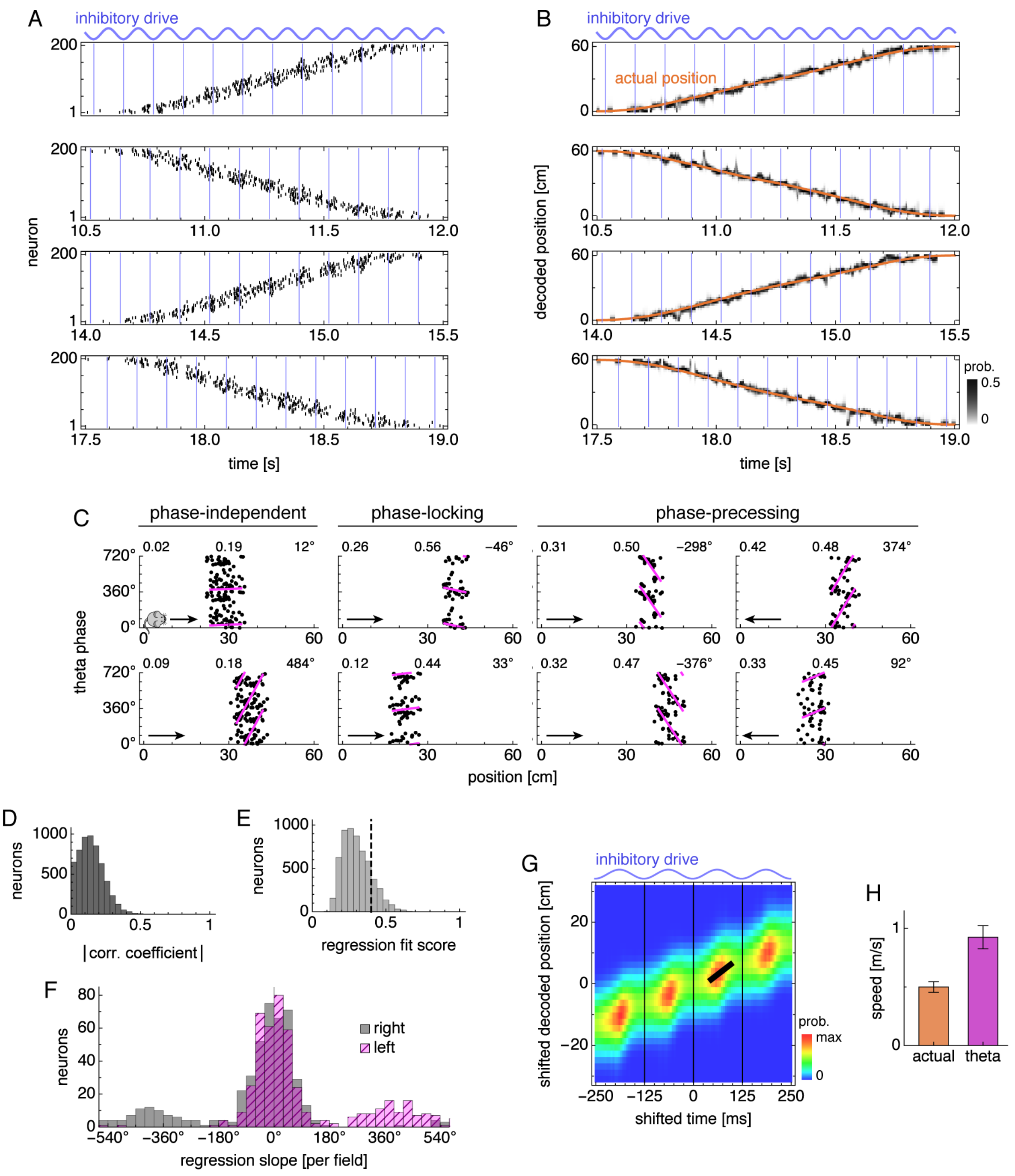


**Figure 8–figure supplement 3**. Phase precession and theta sequence properties in simulations with periodic boundary conditions and uniform excitatory drive. (**A**) Grid cell spike rasters for four runs. Vertical blue lines indicate theta cycle boundaries. (**B**) Decoded position corresponding to **A**. (**C**) Relationship between animal position and theta phase for representative neurons. Dots represent spikes during runs in the directions indicated by arrows. Lines indicate fit by circular-linear regression. Numbers in each panel from top left to top right indicate magnitudes of correlation coefficients, regression fit scores, and regression slopes. (**D–F**) Data across all replicate simulations. (**D**) Magnitudes of circular-linear correlation coefficients. Mean ± s.d.: $0.14 \pm 0.09$. (**E**) Fit scores for circular-linear regression. (**F**) Regression slopes for neurons with fit score $>0.4$. (**G**) Decoded position averaged over theta cycles. Thick black line fit to theta sequence. (**H**) Actual run speed $0.50 \pm 0.05$ m/s (mean ± s.d. over time) and theta sequence speed $0.90 \pm 0.11$ m/s (mean ± s.d. over replicate simulations).
