## Supplementary material for "Replay as wavefronts and theta sequences as bump oscillations in a grid cell attractor network": Captions for videos

### Video captions for "Replay as wavefronts and theta sequences as bump oscillations in a grid cell attractor network"

(Dated: October 25, 2019)

**Video 1.** Neural activity during runs, idle periods, and allocentric corrections. Left, position of the animal (black square) along a 1D track. Right, neural activity of the E population. Each pixel is a neuron, with black corresponding to current spikes and lightest gray corresponding to spikes 40 ms ago. Red circles indicate regions of recording.

**Figure 1–video 1.** Emergence of a grid-like pattern on the neural sheet from randomly initialized membrane potentials. Neural activities of the W and inhibitory populations over the first 240 ms of simulation setup. Each pixel is a neuron, with black corresponding to current spikes and lightest gray corresponding to spikes 40 ms ago. Defects in the grid are then removed as described in the **Model details** subsection of **Appendix A** before the main simulation begins.

**Figure 1–video 2.** Path integration over an open field trajectory produces 2D grid cells. Left, neural activity of the W population. Each pixel is a neuron, with black corresponding to current spikes and lightest gray corresponding to spikes 40 ms ago. Red circle indicates the location of a single recorded neuron. Right, open field trajectory in an enclosure of diameter 1.8 m with a black square indicating the current animal position. Red dots indicate animal positions during spikes recorded from the neuron.

**Figure 3–video 1.** Neural activity during runs and idle periods. Left, position of the animal (black square) along a 1D track. Right, neural activity of the E population. Each pixel is a neuron, with black corresponding to current spikes and lightest gray corresponding to spikes 40 ms ago. Red circles indicate regions of recording.

**Figure 4–video 1.** Phase precession in a simplified conceptual model. An idealized attractor bump, with constant speed and oscillating size, passes through a recorded grid cell multiple times with different initial theta phases. This represents an animal running multiple laps in the same direction at constant speed. Top, the recorded neuron is at location 0 and fires a spike (black dot) whenever it is contained within the bump (gray area), subject to a 40 ms refractory period. Bottom, relationship between theta phase and time for spikes accumulated across multiple laps. Theta phase generally decreases with time within in the grid field. Since the simulated animal moves at constant speed, time is proportional to position, so theta phase decreases with position within in the grid field.

**Figure 4–video 2.** Average attractor bump shape as a function of theta phase. Bump activity is averaged over multiple bumps and theta cycles. Darker areas indicate higher activity, scaled separately for each theta phase. The constant velocity component of bump motion is removed.

**Figure 8–video 1.** Replays preferentially radiate from attractor bumps encoding the current animal position. The activity snapshot in **Figure 8**A corresponds to $t = 41.14\,\mathrm{s}$. Left, position of the animal (black square) at 0 cm along a 1D track, whose positions are colored with a gradient. Right, neural activity of the E population superimposed upon a spatial firing map. Each neuron is colored according the position at which it attains maximum firing rate. Each black pixel represents a spike within the last 5 ms. A wavefront travels through the red recording region located at the top right of the neural sheet. It emanates from an attractor bump sustained by yellow neurons that represent the current position of 0 cm, and it rapidly sweeps through neurons representing progressively farther track positions.

**Figure 8–video 2.** Neural activity during runs, idle periods, and allocentric corrections for a simulation with periodic boundary conditions and uniform excitatory drive. Left, position of the animal (black square) along a 1D track. Right, neural activity of the E population. Each pixel is a neuron, with black corresponding to current spikes and lightest gray corresponding to spikes 40 ms ago. Red circles indicate regions of recording.

**Figure B1–video 1.** Neural activity during runs and idle periods during a simulation without allocentric corrections. Left, position of the animal (black square) along a 1D track. Right, neural activity of the E population. Each pixel is a neuron, with black corresponding to current spikes and lightest gray corresponding to spikes 40 ms ago. Red circles indicate regions of recording.
